# Traumatic Brain Injury Exacerbates Alcohol Consumption and Neuroinflammation with Decline in Cognition and Cholinergic Activity

**DOI:** 10.1101/2024.09.21.614247

**Authors:** Himanshu Gangal, Jaclyn Iannucci, Yufei Huang, Ruifeng Chen, William Purvines, W. Taylor Davis, Arian Rivera, Giles Johnson, Xueyi Xie, Sanjib Mukherjee, Valerie Vierkant, Kaley Mims, Katherine O’Neill, Xuehua Wang, Lee A. Shapiro, Jun Wang

## Abstract

Traumatic brain injury (TBI) is a global health challenge, responsible for 30% of injury-related deaths and significantly contributing to disability. Annually, over 50 million TBIs occur worldwide, with most adult patients at emergency departments showing alcohol in their system. TBI is also a known risk factor for alcohol abuse, yet its interaction with alcohol consumption remains poorly understood. In this study, we demonstrate that the fluid percussion injury (FPI) model of TBI in mice significantly increases alcohol consumption and impairs cognitive function. At cellular levels, FPI markedly reduced the number and activity of striatal cholinergic interneurons (CINs) while increasing microglial cells. Notably, depleting microglial cells provided neuroprotection, mitigating cholinergic loss and enhancing cholinergic activity. These findings suggest that TBI may promote alcohol consumption and impair cognitive abilities through microglia activation and consequently reduced cholinergic function. Our research provides critical insights into the mechanisms linking TBI with increased alcohol use and cognitive deficits, potentially guiding future therapeutic strategies.

## Introduction

Traumatic brain injury (TBI) affects over 50 million people worldwide each year and is a leading cause of death and disability, responsible for approximately 30% of all injury-linked deaths (1, 2). Beyond the immediate physical injuries, TBI often results in long-term consequences such as cognitive impairment, depression, and an increased risk of developing neuropsychiatric diseases, including Parkinson’s disease, Alzheimer’s disease, and substance abuse disorders. Notably, alcohol consumption significantly increases the risk of TBI, with many patients being intoxicated at the time of injury (1–7). This interplay between alcohol and TBI complicates recovery, as alcohol use can impede neurological healing, and TBI itself may increase the likelihood of developing alcohol and substance use disorders (6, 8–13). Despite the profound impact of TBI on cognitive and behavioral functions, the specific mechanisms by which TBI influences alcohol-seeking behaviors and cognitive functions remains poorly understood. Addressing this knowledge gap is critical for developing effective interventaions.

Previous research, including our own, has shown that alterations in striatal activity, particularly in the dorsomedial striatum (DMS), play a key role in regulating alcohol-seeking behaviors and cognitive flexibility (14–16). Specifically, changes in the activity of medium spiny neurons (MSNs) in the DMS and disrupted cortical inputs to these neurons have been linked to increased alcohol consumption and self-administration (15, 16). Additionally, cholinergic interneurons (CINs) in the DMS, although they make up less than 5% of the striatal neuron population, are crucial for regulating drug-induced deficits in cognitive flexibility (14, 17). However, the impact of TBI on these CINs remains largely unexplored. The fluid percussion injury (FPI) model of TBI has been shown to induce significant neuroinflammation in the brain, including the striatum, potentially disrupting striatal circuit activity and leading to cognitive deficits (18–23). Concurrent alcohol exposure further exacerbates this pro-inflammatory state (24), further complicating TBI recovery. CINs, through their release of acetylcholine, modulate both striatal activity and microglia function (25). Microglia activation and chronic neuroinflammation have been linked to cholinergic dysfunction (26–29). However, the specific effects of neuroinflammation on CIN activity following TBI are not well understood.

In this study, we investigated how TBI affects alcohol intake and cognitive function. Our findings indicate that FPI increases alcohol preference and is associated with heightened neuroinflammation and cognitive deficits. FPI also induced a hypo-cholinergic state in the striatum, likely contributing to the observed cognitive deficits. Importantly, depleting microglia using PLX 5622 enhanced cholinergic activity and improved cognitive outcomes. These results highlight striatal neuroinflammation and cholinergic systems as potential therapeutic targets for mitigating the effects of TBI and reducing alcohol intake.

## Results

### FPI increases voluntary alcohol intake and preference

To investigate the impact of FPI on alcohol consumption, male C57BL/6 mice were initially trained to consume 20% alcohol for six weeks using the intermittent-access two-bottle-choice drinking procedure (30–34). Subsequently, animals were assigned to either a sham or FPI group. Mice in the FPI cohort underwent a craniotomy followed by a lateral FPI, while the sham group received only a sterile craniotomy. After surgery, all animals resumed alcohol consumption (Fig. 1a). We found that the sham group showed no significant change in alcohol intake one week (W1) or five weeks (W5) after the procedure (Fig. 1b; *F*(_2,12_) = 1.51; *q* = 0.58, *p* > 0.05 (W1); *q* = 2.35, *p* > 0.05 (W5)). In contrast, the FPI group displayed a marked increase in alcohol consumption both at W1 and W5 post-FPI (Fig. 1e; *F*(_2,20_) = 11.25; *q* = 3.72, *p* < 0.05 (W1); *q* = 6.69, *p* < 0.001 (W5)), indicating that FPI significantly enhanced voluntary alcohol intake in mice.

**Figure 1.**
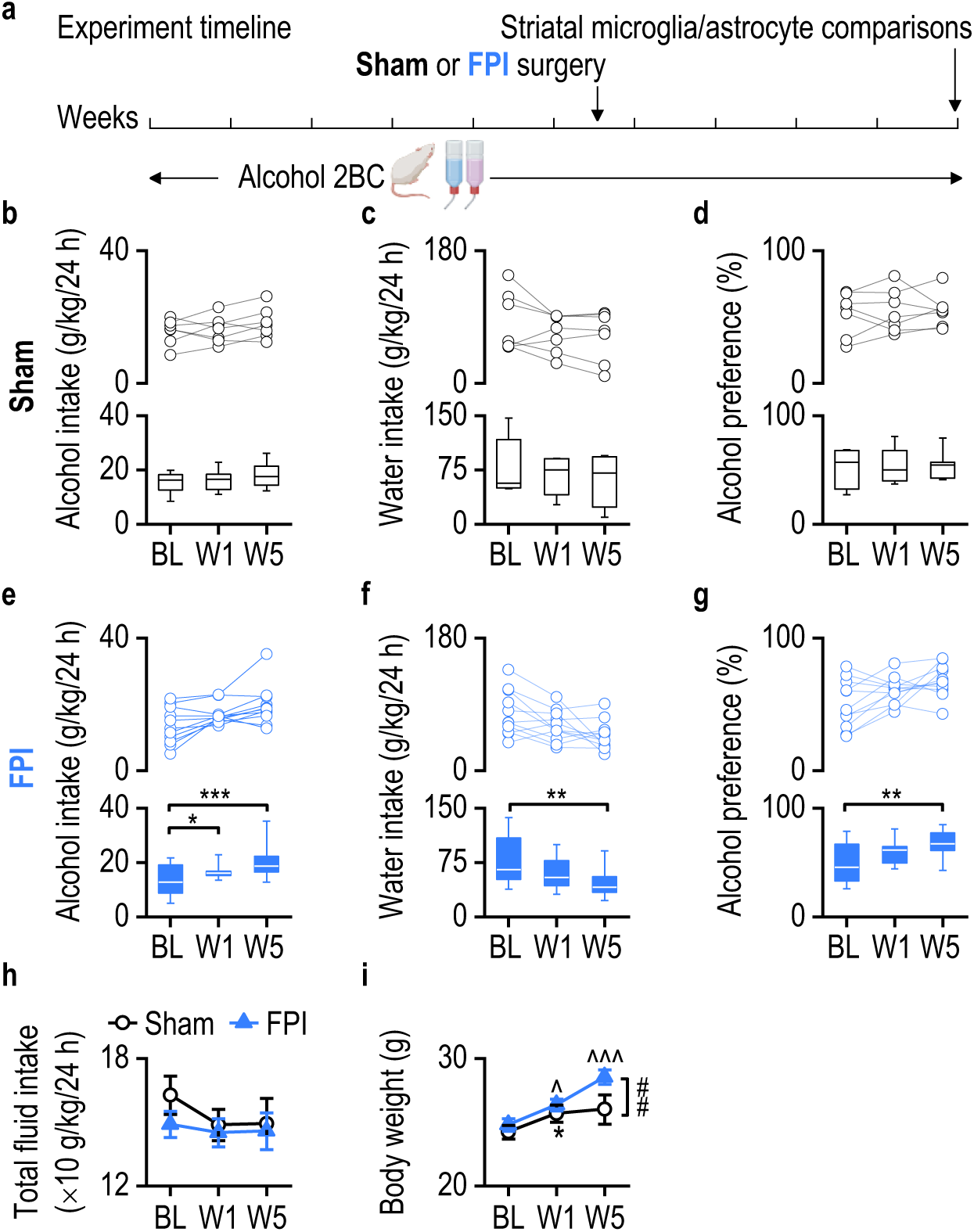
FPI increases alcohol preference in male mice. **a,** Wild-type male C57/BL6 mice were trained to consume 20% alcohol using the intermittent-access two-bottle choice drinking protocol for six weeks, followed by either FPI (blue) or sham (black) surgery. **b,** Alcohol intake did not change in the sham group during weeks 1 (W1) and 5 (W5) post-surgery. **c,** Water intake remained stable post-surgery in the sham group. **d,** Alcohol preference remained unchanged in the sham group post-surgery. **e,** FPI significantly heightened alcohol intake at W1 and W5 post-surgery. **f**, A significant decline in water intake was observed at W1 and W5 post-surgery in the FPI group. **g,** FPI led to a marked increase in alcohol preference at W1 and W5 post-treatment. **h,** Total fluid intake was consistent for both groups before and after their respective surgeries. **i,** FPI, but not sham surgery, led to an increase in body weight at W5 post-surgery. \**p* < 0.05, \*\**p* < 0.01, \*\*\**p* < 0.001; ^*p* < 0.05, ^^^*p* < 0.001 versus baseline (BL) in the same group; ^##^*p* < 0.01 versus sham. One-way RM ANOVA followed by Tukey post-hoc test (b-g), Two-way RM ANOVA followed by Tukey post-hoc test (h, i). n = 7 (Sham), 11 (FPI).

Water intake was also monitored, revealing no significant changes in the sham group during W1 and W5 post-surgery (Fig. 1c; *F*(_2,12_) = 2.53, *p* > 0.05). However, the FPI group showed a notable reduction in water intake at W5 post-FPI compared to baseline levels (Fig. 1f; *F*(_2,20_) = 6.75, *p* < 0.01 (W5)), suggesting a shift in fluid preference towards alcohol. Indeed, while the sham group did not demonstrate significant changes in alcohol preference post-surgery (Fig. 1d; *F*_(2,12)_ = 0.19, *p* > 0.05), the FPI group exhibited a significant increase in alcohol preference at W5 post-FPI relative to baseline (Fig. 1g; *F*_(2,20)_ = 6.37, *p* < 0.01 (W5)). Total fluid intake (alcohol and water combined) remained comparable between groups (Fig. 1h; *F*_(1,16)_ = 0.51, *p* > 0.05) and throughout the study period (Fig. 1h; *F*_(2,32)_ = 1.48, *p* > 0.05).

Body weight was measured before and after surgery in both groups. The sham group exhibited a slight increase in body weight at W1 post-surgery (Fig. 1i; *q* = 3.59, *p* < 0.05). However, the FPI group displayed a more substantial increase in body weight both at W1 and W5 post-injury (Fig. 1i; *q* = 3.99, *p* < 0.05 (W1 versus BL); *q* = 9.51, *p* < 0.001 (W5 versus BL); *q* = 5.52, *p* < 0.01 (W5 versus W1)), with significantly higher body weights observed at W5 compared to the sham group (Fig. 1i; *q* = 3.91, *p* < 0.01 (W5)). This continuous weight gain in the FPI group, correlated with increased alcohol access, aligns with studies showing weight gain following TBI (35–38).

These results collectively suggest that FPI not only increases alcohol consumption and preference in mice but also affects other physiological parameters such as body weight and water intake.

### FPI increases microglial activity in the ipsilateral and contralateral striatum

Given the critical role of microglia in neuroinflammation following TBI or alcohol exposure (39), we next investigated how TBI influences microglial activity in alcohol-drinking mice. To achieve this, we assessed Iba-1 labeled microglial cells in the striatum at W5 following either sham or FPI treatment (Fig. 2a, b). The analysis revealed a significant increase in microglial optical density in the ipsilateral striatum following FPI as compared to sham controls (Fig. 2c; *F*_(1,66)_ = 23.70, *p* < 0.001). A post-hoc Tukey test confirmed significant elevations in all medial subregions of the ipsilateral striatum in the FPI-treated mice compared to the sham group (Fig. 2c; dorsal medial, *q* = 3.58, *p* < 0.05; central medial, *q* = 3.08, *p* < 0.05; ventral medial, *q* = 5.18, *p* < 0.001). However, increases in the optical density of lateral subregions in the ipsilateral striatum, though apparent, were not statistically significant (Fig. 2c; dorsal lateral, *q* = 1.76, *p* = 0.22; central lateral, *q* = 1.55, *p* = 0.28; ventral lateral, *q* = 1.72, *p* = 0.23). Collectively, these changes resulted in a significant overall increase in the total microglial optical density (Fig. 2d; *t*_(11)_ = -2.38, *p* < 0.05) and in the total number of microglial cells (Fig. 2e; *t*_(11)_ = -2.22, *p* < 0.05) within the ipsilateral striatum.

**Figure 2.**
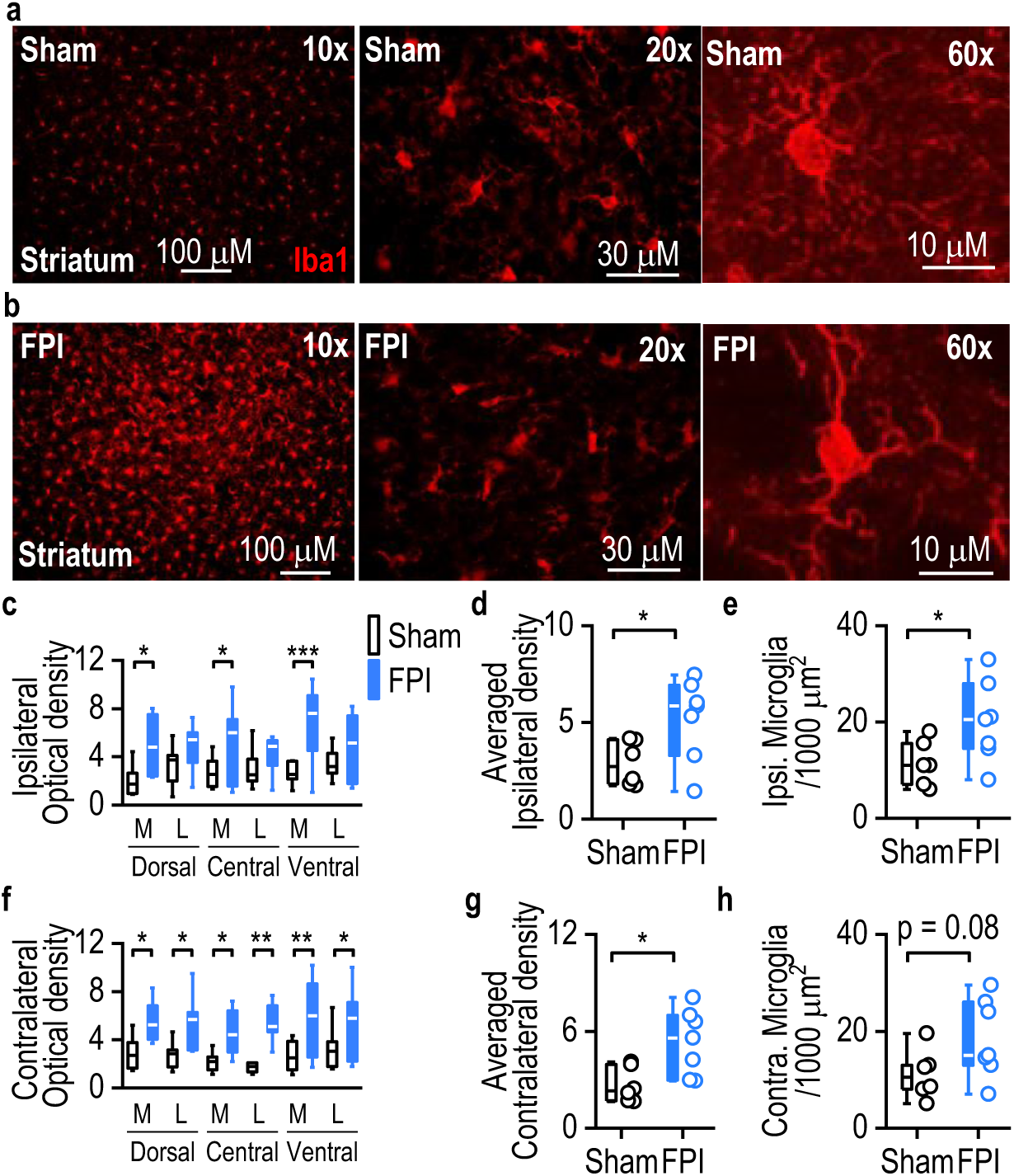
FPI increases microglial levels in the striatum. **a, b,** Confocal micrographs display Iba-1-stained coronal striatum sections from sham plus alcohol (top, a) and FPI plus alcohol (bottom, b) groups. **c-e,** In the ipsilateral striatum, FPI significantly increased the microglial optical density in the medial regions (c) and across the entire ipsilateral striatum (d). Additionally, there was an increase in the total number of microglial cells on the ipsilateral side (e). **f-h,** In the contralateral striatum, FPI significantly enhanced the optical density across all subregions (f) and in the entirety of the contralateral striatum (g). There was a marginal increase in the number of microglial cells in the contralateral striatum (h). Iba-1, Ionized calcium-binding adaptor molecule 1; M, Medial; L, Lateral. \**p* < 0.05, \*\**p* < 0.01, \*\*\**p* < 0.001. Two-way RM ANOVA followed by Tukey post-hoc test (c, f), unpaired t-test (d, e, g, h). n = 6 (a-h, Sham), 7 (a-h, FPI).

Interestingly, we also observed a significant enhancement in the microglial optical density after FPI across all subregions of the contralateral striatum (Fig. 2f; *F*_(1,66)_ = 37.40, *p* < 0.001; dorsal medial, *q* = 3.29, *p* < 0.05; dorsal lateral, *q* = 3.35, *p* < 0.05; central medial, *q* = 3.13, *p* < 0.05; central lateral, *q* = 4.34, *p* < 0.01; ventral medial, *q* = 4.16, *p* < 0.01; ventral lateral, *q* = 2.92, *p* < 0.05). This increase was consistent throughout the contralateral striatum (Fig. 2g; *t*_(11)_ = -2.89, *p* < 0.05). A marginal increase was also noted in the number of microglial cells in the contralateral striatum after FPI (Fig. 2h; *t*_(11)_ = -1.91, *p* = 0.08). These results indicate that FPI induces an increase in microglial activity in both the ipsilateral and contralateral striatum, suggesting a widespread neuroinflammatory response.

### FPI does not alter the number of astrocytes in the striatum

While microglia activation was evidence following FPI, it is also important to consider the potential effects on other glial cells, such as astrocytes (40). To asses whether FPI affects astrocyte numbers in the striatum of alcohol-drinking mice, we analyzed GFAP-labeled astrocytes in both the ipsilateral and contralateral striatum (Fig. 3a, d). Our analysis revealed no significant changes in the optical density of astrocytes within the subregions of the ipsilateral striatum between the FPI and sham groups (Fig. 3b; *F*_(1,66)_ = 1.52, *p* > 0.05), as well as in the overall ipsilateral striatum (Fig. 3c; *t*_(11)_ = - 0.72, *p* > 0.05). Similarly, there were no significant differences in astrocyte optical density between the FPI and sham groups in the subregions of the contralateral striatum (Fig. 3e; *F*_(1,66)_ = 0.87, *p* > 0.05), or in the overall contralateral striatum (Fig. 3f; *t*_(11)_ = 0.01, *p* > 0.05). These findings collectively suggest that FPI does not significantly affect the number of astrocytes in either the ipsilateral or contralateral striatum in alcohol-drinking mice.

**Figure 3.**
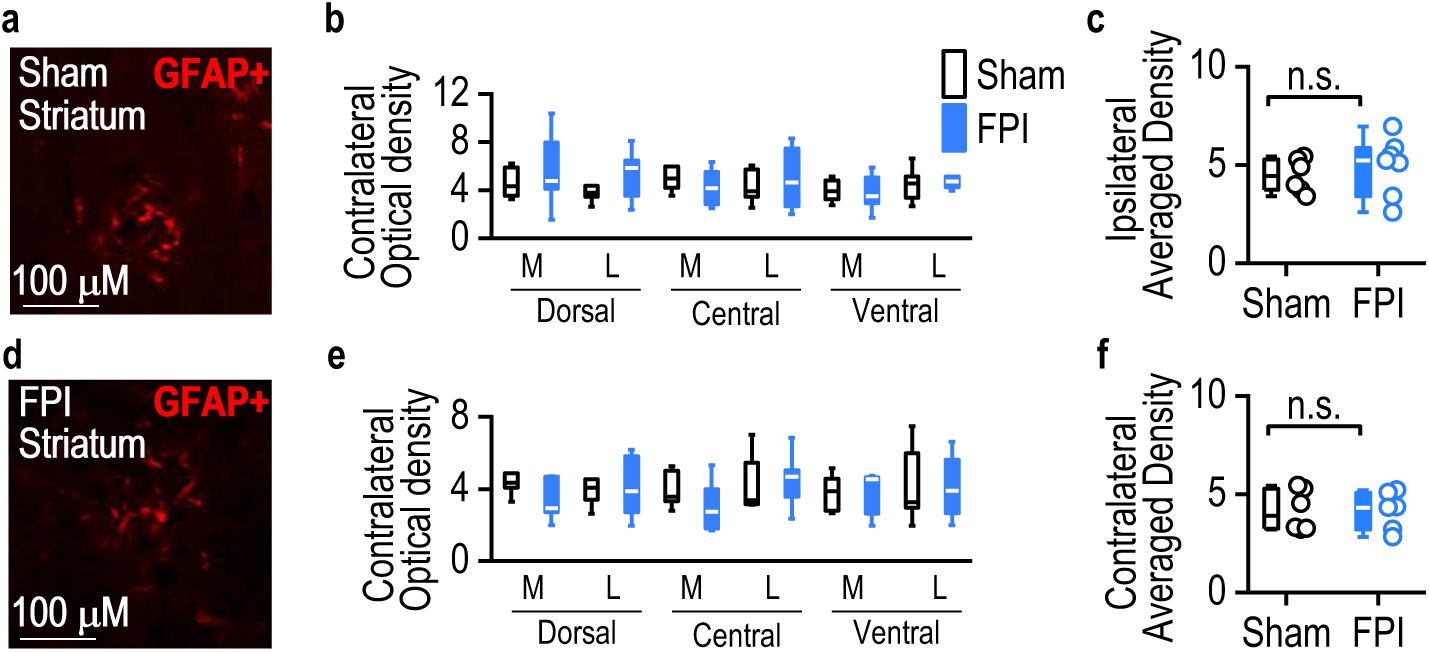
FPI does not alter astrocytes in the striatum. **a,** Confocal micrographs show GFAP-stained coronal striatum sections from the sham plus alcohol group. **b-c,** In the ipsilateral striatum, FPI did not significantly alter astrocyte optical densities in striatal subregions (b) or the average density of astrocytes in the entire ipsilateral striatum (c). **d,** Confocal micrographs of GFAP-stained coronal striatum sections from the FPI plus alcohol group. **e-f,** In the contralateral striatum, FPI did not significantly alter astrocyte optical densities in striatal subregions € or the average density of astrocytes in the entire contralateral striatum (f). n.s. (not significant; c, f). Two-way RM ANOVA followed by Tukey post-hoc test (b, e), unpaired t-test (c, f). n = 6 (a-f, Sham), 7 (a-f, FPI).

### FPI causes cognitive but not motor impairment

While FPI alters the number of microglia, but not astrocytes in the striatum of alcohol-drinking mice, its impact on cognitive function and locomotor activity remains evident, Given that TBI and chronic alcohol intake both affect cognitive function (41, 42), we assessed the combined effect of FPI and alcohol intake on locomotion, exploratory behavior, and cognitive function using open-field and novel object recognition tests (NORT) four weeks post-surgery (Fig. 4a). The open-field test revealed no significant differences in total distance traveled (Fig. 4b, c; *t*_(16)_ = -0.60; *p* > 0.05) or locomotion velocity (Fig. 4d; *t*_(16)_ = -0.18; *p* > 0.05) between the FPI and sham groups, indicating similar levels of locomotor activity. The amount of time spent in the periphery of the arena was also comparable between these two groups (Fig. 4e; *t*_(16)_ = 0.13; *p* > 0.05), suggesting no differences in anxiety response to a novel environment between them.

**Figure 4.**
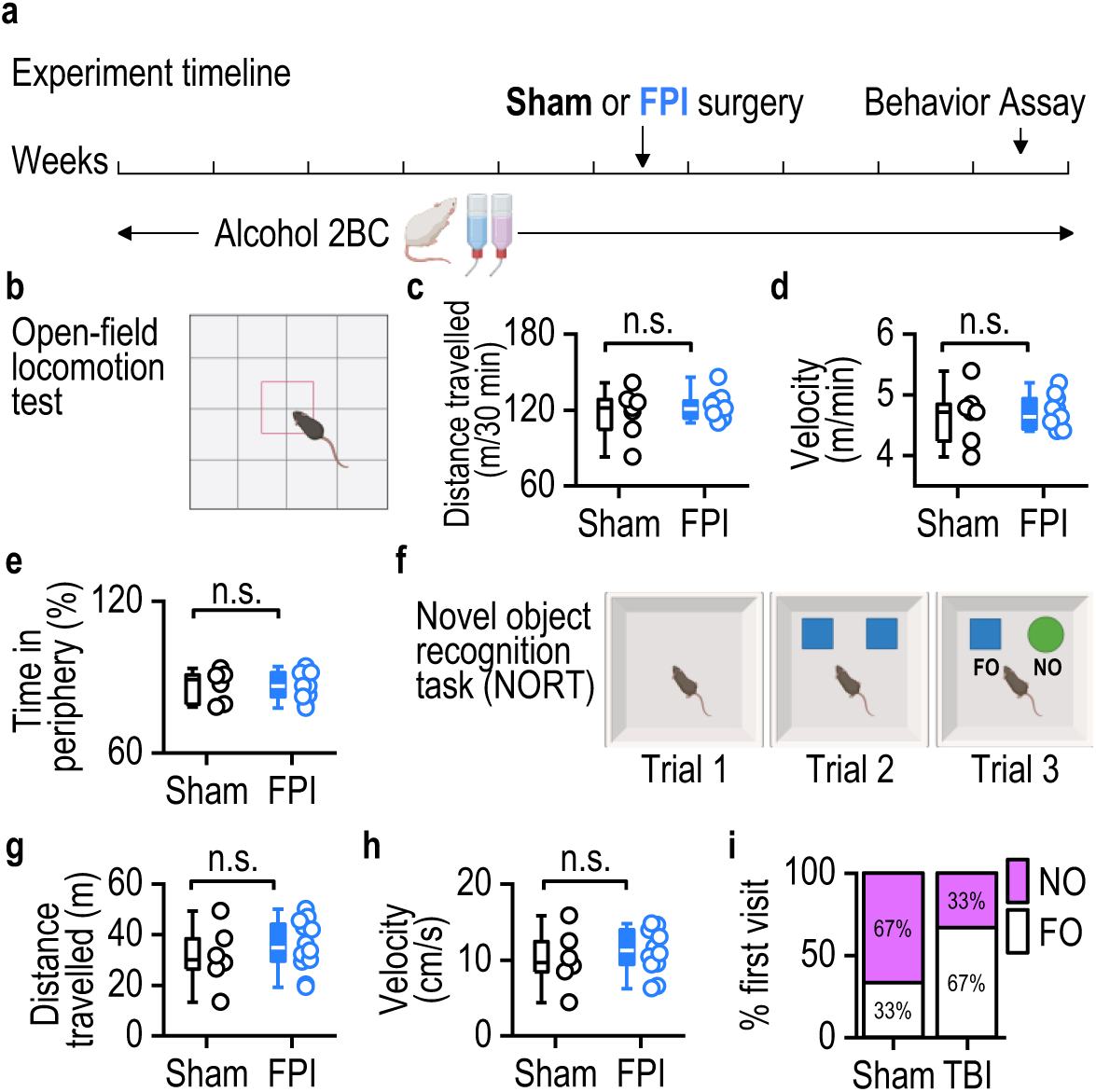
FPI causes cognitive but not motor deficits in mice with an alcohol-drinking history. **a,** Experiment timeline. **b-e,** No significant changes were observed in total distance traveled I, movement velocity (d), or time spent in the periphery of the open field chamber I after FPI. **f-h,** In the NORT, FPI did not affect total distance traveled (g) or velocity of movement (h). **I,** FPI-treated mice’s first approach to the novel object was less than sham controls, indicating a cognitive deficit. N.s. (not significant; c, d, e, g, h). \**p* < 0.05 (i). unpaired t-test (c, d, e, g, h, i). n = 6 (Sham), 12 (FPI).

Cognitive function was assessed using NORT, a task measuring reference memory (Fig. 4f). Despite similar levels of physical activity in the arena, as indicated by no significant differences in distance traveled (Fig. 4g; *t*_(16)_ = 0.8103; *p* > 0.05) or velocity (Fig. 4h; *t*_(16)_ = 0.7093; *p* > 0.05) between groups, the FPI group exhibited significant cognitive deficits. Significantly less percentage of FPI mice first approached the novel object as compared to the sham controls during the novel object recognition task (Fig. 4i; *p* < 0.05). Therefore, while the FPI and alcohol exposure did not affect motor capabilities, they did lead to cognitive impairments, as evidenced by decreased novelty preference in NORT.

### FPI reduces cholinergic activity in the DMS

Despite the preservation of motor capabilities, FPI and alcohol exposure impairs cognitive function. Given the association between cognitive deficits and diminished activity in striatal CINs (17, 42), we further investigated the effects of FPI on these neurons using three groups of ChAT-eGFP mice: a naïve group, a group subjected only to FPI, and a group with both alcohol consumption and FPI treatment. Five weeks post-FPI, the animals were euthanized, and their brains were processed for confocal scanning to count the total number of striatal CINs. We observed that both the FPI-only and the alcohol+FPI groups exhibited a lower number of striatal CINs per slice compared to the naïve group (Fig. 5a, b; *t*_(1, 44)_ = -2.98, *p* < 0.01 (FPI); *t*_(1, 48)_ = -2.21, *p* < 0.05 (alcohol+FPI); Supplementary Fig. 1a, b). However, there was no significant difference in CIN counts between the FPI-only and the alcohol+FPI groups (Fig. 5b; *t*_(1, 42)_ = -1.09, *p* > 0.05). These findings suggest that FPI leads to a reduction in the number of striatal CINs, that is not significantly exacerbated by concurrent alcohol exposure.

**Figure 5.**
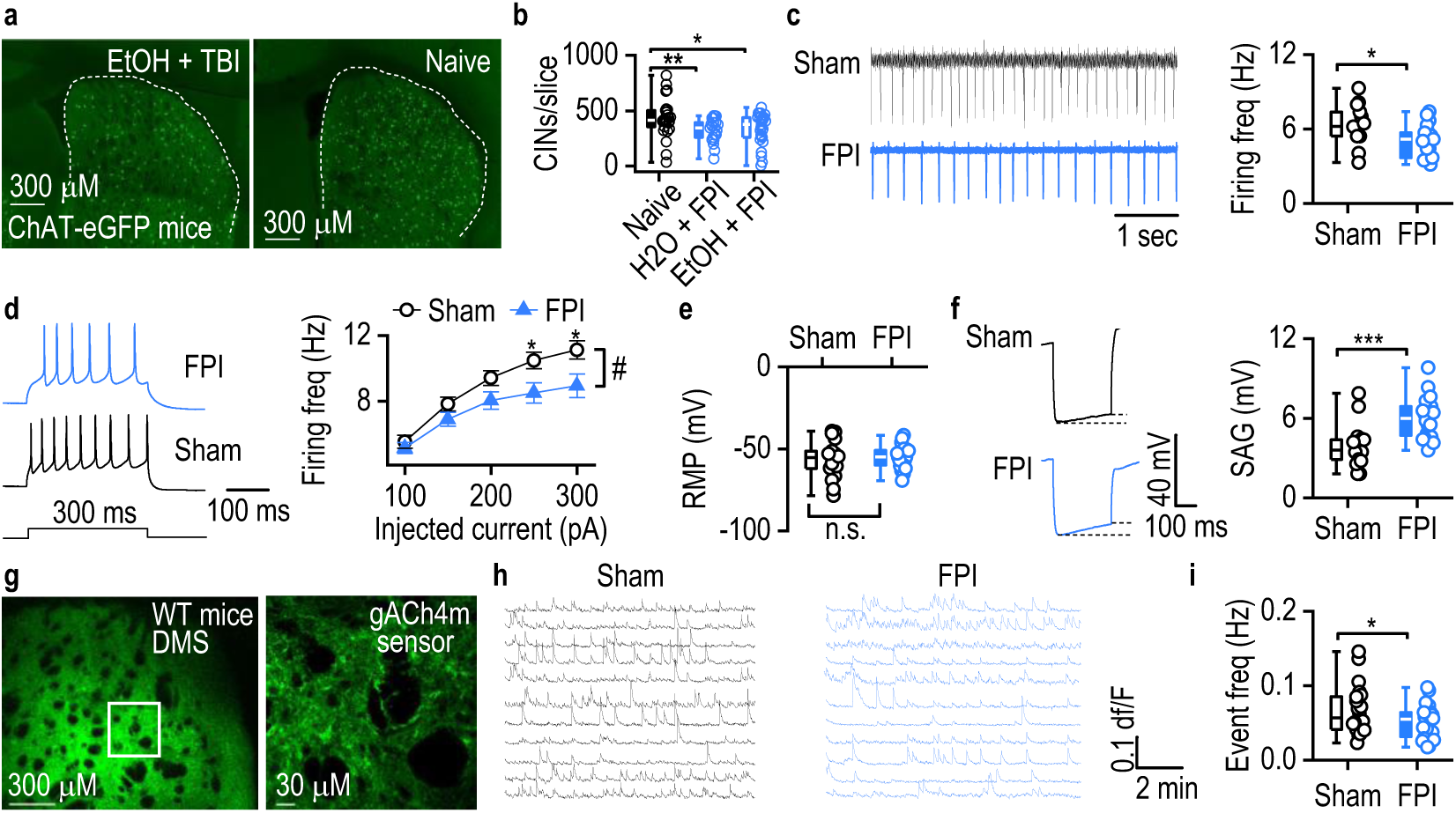
FPI suppresses DMS cholinergic activity. **a,** Confocal micrographs show striatal sections from the EtOH+FPI and naïve groups, with green neurons representing CINs. **b,** FPI significantly decreased the number of CINs per slice in both water and alcohol-exposed groups compared to naïve controls. **c,** FPI surgery reduced spontaneous firing rates of DMS CINs. **d,** FPI reduced the number of evoked action potentials in DMS CINs. **e,** FPI did not significantly alter the resting membrane potential of DMS CINs. **f,** FPI significantly increased the SAG in DMS CINs. **g,** Confocal micrographs show acetylcholine sensor expression in the striatum of wild-type mice. **h-i,** FPI reduced the frequency of acetylcholine events in the dorsomedial striatum. n.s. (not significant; e), ^#^*p* = 0.07, \**p* < 0.05, \*\**p* < 0.01, \*\*\**p* < 0.001. Two-way RM ANOVA followed by Tukey post-hoc test (d), unpaired t-test (b, c, e, f, i). n = 26 (slices) and 3 (mice) (26/3, b, Naive), 24/4 (b, H2O+FPI), 24/3 (b, EtOH+FPI). n = 13 (neurons) and 3 (mice) (13/3, c, sham), 17/4 (c, FPI), 17/3 (d, e, f, sham), 28/4 (d, e, FPI), 21/3 (f, FPI), n = 22 (slices) and 4 mice (22/4, i, sham), 25/5 (i, FPI).

Further, we investigated the spontaneous activity of CINs in the DMS. ChAT-eGFP mice were either given FPI or sham surgery and after a week of recovery, DMS slices were prepared for electrophysiology recordings. The CINs were identified by their green fluorescence. In cell-attached recordings, it was evident that the spontaneous firing frequency of DMS CINs was significantly reduced in the FPI group compared to sham controls (Fig. 5c; *t*_(28)_ = 2.44, *p* < 0.05). Additionally, in current-clamp recordings that involved injecting small step currents, the number of evoked action potentials in CINs was lower in the FPI group, though this finding approached but did not reach statistical significance (Fig. 5d; *F*_(1, 43)_ = 3.43, *p* = 0.07). There was no observed change in the resting membrane potential between the two groups (Fig. 5e; *t*_(43)_ = -0.59, *p* > 0.05). Notably, FPI also resulted in a significant increase in the characteristic sag response in DMS CINs, indicative of changes in the hyperpolarization-induced cation current, *I*_h_ (Fig. 5f; *t*_(36)_ = -4.18, *p* < 0.001). These results collectively suggest that FPI reduces cholinergic activity in the DMS, impacting the functionality of CINs.

In voltage-clamp recordings, we observed no significant differences in AMPA-induced currents between the FPI and sham groups (Supplementary Fig. 1c; *t*_(16)_ = -0.73, *p* > 0.05). Similarly, FPI did not affect the frequency (Supplementary Fig. 1d; *t*_(26)_ = 1.37, *p* > 0.05) or amplitude (Supplementary Fig. 1e; *t*_(26)_ = -0.29, *p* > 0.05) of spontaneous inhibitory postsynaptic currents (sIPSCs). Additionally, there were no changes in the frequency (Supplementary Fig. 1f; *t*_(21)_ = -0.42, *p* > 0.05) or amplitude (Supplementary Fig. 1g; *t*_(21)_ = -0.62, *p* > 0.05) of spontaneous excitatory postsynaptic currents (sEPSCs). The lack of changes in sEPSCs or sIPSCs suggests FPI may not affect glutamatergic or GABAergic transmission in the DMS.

Given the observed reduction in striatal CIN activity due to FPI, we next examined whether FPI could also affect acetylcholine release in the DMS. For this purpose, an adeno-associated virus (AAV) encoding a green acetylcholine sensor (gACH4m) was infused in wild-type mice (Fig. 5g). Two weeks after infusion, the animals underwent either FPI or sham surgeries. Following a one-week recovery period, live tissue confocal recordings were performed to assess acetylcholine release in the DMS of both groups (Fig. 5h). The results indicated a significant decrease in the frequency of spontaneous fluorescent events in the FPI group compared to the sham controls (Fig. 5i; *t*_(45)_ = 2.099, *p* < 0.05). These findings suggest that FPI reduces cholinergic activity in the DMS, which may contribute to the observed cognitive deficits following FPI.

### PLX 5622-induced microglia depletion increased striatal acetylcholine activity and improved cognitive function

After demonstrating that FPI combined with alcohol consumption significantly increased striatal microglia and decreased striatal acetylcholine activity, we next investigated whether microglia depletion, achieved through PLX 5622 administration, could influence CIN activity and address the cognitive deficits observed after FPI. Previous findings showed microglial cells in close proximity to cholinergic projection neurons in the basal forebrain (43). Consistently, we discovered that microglia cells are also close to CINs in the DMS (Supplementary Fig. 2a). To deplete microglia, mice received intraperitoneal injections of PLX 5622 (50 mg/kg) twice daily (12-h intervals) for 7 days, while control mice were administered saline (Fig. 6a). Iba-1 staining confirmed that PLX 5622 treatment significantly reduced microglial presence in the brain (Fig. 6b), with a 60% reduction in striatal microglia observed in the PLX-treated mice compared to controls (Fig. 6c; *t*_(16)_ = 4.089, *p* < 0.001).

**Figure 6.**
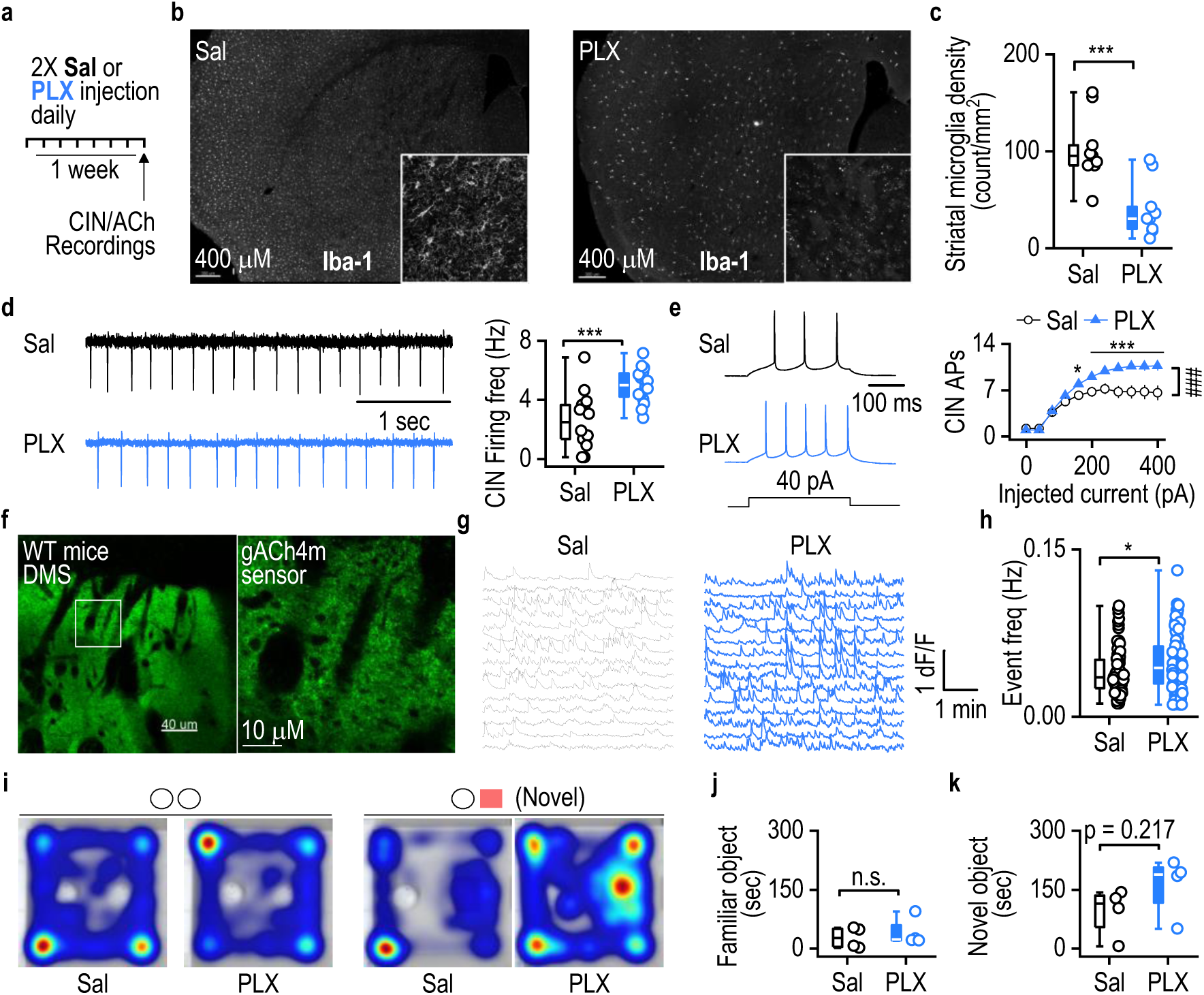
Microglia depletion increases striatal acetylcholine activity. **a,** Mice received injections of either PLX 5622 (50 mg/kg) or saline twice daily for one week. Electrophysiology and confocal recordings were performed one day after the last PLX injection. **b,** Confocal micrographs illustrate the PLX-induced reduction in microglial cells; the right bottom box shows a magnified image from the striatum. **c,** Summarized data showing microglia depletion after PLX 5622 administration. **d,** PLX administration led to an increase in spontaneous firing frequency of DMS CINs. **e,** PLX treatment increased the number of current-evoked action potentials in DMS CINs. **f,** An acetylcholine sensor (gACh4m) was expressed in the striatum of wild-type mice. **g-h,** Live-tissue confocal recordings of striatal acetylcholine (g) revealed increased baseline acetylcholine events in PLX-treated mice compared to saline controls (h). **i,** On the 6^th^ day of PLX administration, mice underwent familiar object training, followed by a NORT on the 7^th^ day. **j,** No difference in the time spent with the familiar object was observed during the NORT between the groups. **k,** Trend of increased visitation of the novel object in PLX-treated mice compared to controls. n.s. (not significant; j), \**p* < 0.05, ^###^*p* < 0.001, \*\*\**p* < 0.001. Two-way RM ANOVA followed by Tukey post-hoc test (e), unpaired t-test (c, d, h, j, k). n = 9 slices from 3 mice (9/3, c, PLX, Sal), n = 14 (neurons) and 4 (mice) (14/4, d, e, Sal), 16/4 (d, e, PLX), n = 91 (ROIs) from 4 (mice) (91/4, h, Sal), 98/4 (h, PLX), n = 4 mice (j, k; Sal, PLX).

Subsequently, we examined CIN activity in ChAT-eGFP mice treated with either PLX 5622 or saline. Slice recordings in DMS CINs, conducted a day after the final PLX 5622 treatment, revealed that the spontaneous firing frequency was significantly higher in the PLX group than in saline controls (Fig. 6d; *t*_(28)_ = 4.184, *p* < 0.001). Similarly, current-clamp recordings showed that the same current injection evoked more action potentials in CINs from PLX-treated mice than in those from saline controls (Fig. 6e; *F*_(1,20)_ = 25.36, *p* < 0.001). We also measured baseline striatal acetylcholine activity using live-tissue confocal imaging in wild-type mice expressed with an acetylcholine sensor in the DMS (Fig. 6f, g). Results from the day following the last PLX 5622 treatment indicated a significantly higher frequency of spontaneous acetylcholine events in the DMS of PLX-treated mice than saline controls (Fig. 6h; *t*_(187)_ = -2.521, *p* < 0.05), though no differences were observed in amplitude or half-width of these events (Amplitude: Supplementary Fig. 2b, t_(16)_ = -1.413, *p* > 0.05; Half-width: Supplementary Fig. 2c, t_(16)_ = 0.0597, *p* > 0.05). These results suggest that microglia depletion enhances CIN activity and acetylcholine release in the DMS.

Striatal CINs and acetylcholine activity are known to positively influence cognitive behaviors. Therefore, we hypothesized that the enhancement of striatal acetylcholine activity by PLX 5622 could improve performance in cognitive tasks such as the novel object recognition test, conducted on the last day of PLX injections. Our observations indicated no significant differences in the time spent near the familiar object between PLX-treated and saline-treated control mice (Fig. 6j, *t*_(6)_ = -0.592, *p* > 0.05). However, there was a notable trend toward increased time spent near the novel object by PLX-treated mice compared to saline controls (Fig. 6k, *t*_(6)_ = -1.378, *p* > 0.05). Collectively, these findings suggest that microglia depletion via PLX 5622 enhances striatal acetylcholine activity and could potentially improve cognitive function.

## Discussion

In this study, we found that FPI-induced TBI significantly increased alcohol preference in mice. When combined with alcohol, FPI also resulted in cognitive impairment and a chronic elevation in the number of striatal microglial cells but not of astrocytes. Furthermore, FPI reduced the number and spontaneous activity of CINs, as well as acetylcholine release in the striatum. Interestingly, depleting microglia was found to enhance cholinergic activity in FPI-untreated mice. These results align with both preclinical and clinical studies that report increased alcohol consumption following TBI (44–47). Given the association between TBI and heightened risks of binge drinking and alcohol use disorders, these anatomical and functional changes in the striatum post-FPI are particularly significant.

Previous studies have indicated that neuroinflammation, specifically the activity of microglial cells, may play a crucial role in increasing alcohol consumption (46). Supporting this notion, the administration of minocycline, which targets microglial activation, was shown to prevent the rise in alcohol consumption induced by TBI in juvenile mice subjected to a mild closed-head FPI (46). Additionally, direct alcohol injection into the ventral striatum triggered microglial activation, which was mitigated by minocycline (46). In rat models of lateral FPI combined with post-injury alcohol consumption, neuroinflammation was observed in the peri-injury cortex at 2-3 weeks post-FPI, correlating with increased alcohol intake and cognitive deficits (8, 48). Moreover, acute alcohol intoxication has been shown to prolong neuroinflammation in rats following FPI (49). Conversely, a closed-head TBI model demonstrated increased astrocytosis as early as 72 h post-injury, persisting for at least 7 days, including in the nucleus accumbens. This was linked to reduced alcohol consumption immediately after the injury (50), further highlighting the complex relationship between microglia-mediated neuroinflammation and elevated alcohol preference after injury. It is pertinent to note that our examination of astrocytes was at a much more chronic timepoint post-TBI.

In our current investigation of the dorsal striatum following lateral FPI in young adult mice, we observed chronic microglial activation within our voluntary alcohol consumption paradigm. While the ventral striatum is known to play a central role in driving reinforcement behaviors (51), our studies, along with others, have identified the involvement of dorsal striatal circuitry in promoting alcohol-seeking behavior (15, 16). This suggests that microglial activation in both the dorsal and ventral striatum may contribute to the mechanisms underlying post-TBI alcohol-seeking behaviors.

To investigate the neural mechanisms underlying the cognitive decline observed after FPI, we conducted electrophysiological recordings of CINs in the dorsal striatum, given their critical role in regulating cognitive flexibility (14, 17). The observed reduction in the activity of these CINs is particularly noteworthy, as it correlates with increased alcohol consumption and preference. Similar alterations in CINs have been previously implicated in the compulsivity of drug use (52), suggesting that TBI may directly affect striatal circuitry, thereby enhancing the signals that drive alcohol preference. Given the established effects of neuroinflammation on neural circuitry, it is plausible that the impacts on these neurons stem from FPI-induced neuroinflammation. The chronic elevation of microglia cells in this region following FPI and alcohol consumption indicates that alcohol may be a driving factor for sustained chronic microglial activation, as observed in models involving alcohol but not TBI (53). Considering the involvement of microglial cells in synaptic maintenance, pruning, and plasticity, further investigation into the role of these activated microglia in the physiological changes of CINs would be valuable.

The deficits noted in striatal cholinergic interneurons may underpin the cognitive impairments observed in the NORT post-FPI. NORT, a test that depends on the functionality of multiple brain regions including the hippocampus, perirhinal cortex, and the mFPC, is sensitive to disruptions in object recognition linked to the striatum (54–60). In rodent models of schizophrenia, object recognition deficits were mediated by reductions in VGLUT1 across the mPFC, striatum, and hippocampus (61). Moreover, altered corticostriatal communication was found to underlie recognition deficits, characterized by reduced tonic firing of neurons in the nucleus accumbens and desynchronization of mPFC and nucleus accumbens neurons (62). Although the nucleus accumbens is part of the ventral striatum and our findings focus on neuronal activity in the DMS, these results support the notion that disrupted regional connectivity underlies the object recognition deficits observed following alcohol consumption and FPI comorbidity. Future studies should consider examining the changes in regional connectivity resulting from alcohol consumption and TBI, to further elucidate these relationships.

Another contributing factor to the alterations observed in NORT could be linked to disrupted cholinergic signaling, which has been associated with object recognition deficits (63). Striatal CINs are crucial for various cognitive processes (63–65) and particularly important for the acquisition of object recognition memory (66). Thus, the FPI and alcohol-induced changes in these interneurons in the striatum likely underpin the cognitive impairments we identified.

In conclusion, this investigation highlights significant connections between TBI, alcohol consumption, and their consequent impact on the brain. Although beyond the scope of the current study, it would be intriguing to explore whether the observed increase in alcohol intake correlates with a decline in glucose metabolism. Additionally, determining whether an increase in body weight is driven by food intake or the caloric contributions of alcohol could offer valuable insights for future research. The elevated presence of microglial cells in the striatum and alterations in striatal CINs underscore how brain inflammation might enhance alcohol preference following an FPI. These changes not only relate to cognitive impairments but also suggest that modifications in brain region connectivity could contribute to these challenges. Furthermore, disruptions in cholinergic signaling within the striatum could elucidate the cognitive difficulties observed following FPI. These findings may provide clues about the pathogenic mechanisms that elevate the risk for alcohol and substance use disorders after TBI.

## Materials and Methods

### Sex as a biological variable

Male C57BL/6 mice were initially group housed (n = 4 per cage) at the Texas A&M University Health Science Center animal facility, maintained in a temperature (72 degrees fahrenheit)- and humidity (54 percent)-controlled environment, with ad libitum access to food and water. Post sham/FPI treatment, mice were single housed. Male mice were used in this study because the incidence of TBI in young adults is known to be more prominent in males, however the findings are expected to be relevant for female mice as well. Detailed descriptions of the experimental treatments are provided below.

### Intermittent-Access to 20% Alcohol 2-Bottle-Choice Drinking Procedure

Mice were trained to intermittently consume 20% alcohol for six weeks, following a method previously described (17, 30, 33, 42, 67–70). Alcohol consumption was monitored before the mice were divided into the FPI and sham treatment groups, ensuring equal distribution of drinking tendencies between groups prior to intervention. After sham or FPI induction, intermittent access to alcohol continued for an additional five weeks, during which consumption was monitored. Behavioral testing was performed over the following week before the mice were euthanized.

### Fluid Percussion Injury

After six weeks of alcohol drinking, mice were anesthetized and subjected to lateral FPI as previously performed (18, 71, 72). Specifically, the animals were anesthetized, placed in a stereotaxic instrument, and underwent a sterile craniotomy over the left parietal cortex. Mice in the sham group were connected to the FPI apparatus without receiving an injury, whereas the FPI group received a lateral FPI. The injury involved a 12-16 ms pressure pulse delivered at ∼1.4-2.6 atm from the FPI apparatus (Custom Design & Fabrication, Richmond, VA). Following injury, the incision sites were closed with sterile sutures and mice were returned to their home cages for recovery.

### Novel Object Recognition Test (NORT)

The NORT assesses memory and exploratory behavior in rodents (55). This test included three successive trials with 1-h inter-trial interval (ITI). Each trial was conducted in a white plexiglass open field box (60 cm x 45 cm). In the first trial, mice were allowed to explore the empty box for 1 hour to acclimate and assess general locomotor ability. In the second trial (training trial), two similar objects were placed in the open field box. In the third trial, one object from the training trial was replaced with a new object (termed the novel object, NO), while the other object (termed the familiar object, FO) remained. All trials were videotaped for both automatic and manual scoring. Locomotor activity in the first trial was assessed using a Kinder Scientific SmartFrame Open Field System, focusing on distance traveled, locomotion velocity, and time spent in the periphery. During the second and third trials, object exploration was scored based on whether the mouse’s nose contacted an object, including metrics such as the first object visited, latency to the first object, and latency to NO.

### Tissue Processing and Histology

Upon completion of behavioral testing, mice were euthanized via transcardial perfusion as previously described (17, 72–75). Following perfusion and a 24-hour post-fixation period in situ, brains were extracted, post-fixed for an additional 24-48 h in 4% paraformaldehyde (PFA), and subsequently transferred to phosphate-buffered saline (PBS). Brains were sectioned in the coronal plane using a Vibratome (LEICA VT 1200) at 50-µm intervals. Serial sections were collected sequentially in 12-well plates and stored at 4°C in a 0.01% PBS solution. Astrocytes were stained using an anti-GFAP-Cy3 conjugated antibody (Sigma), and microglia were labeled with primary anti-Iba-1 (WAKO labs) and secondary goat anti-rabbit IgG (Invitrogen). After staining, sections were mounted onto glass slides, covered with VECTASHIELD (Vector Labs, Burlingame, CA), and sealed with nail polish to secure the coverslips. Striatal slices were imaged using confocal microscopy.

Images focusing on the dorsal striatum (bregma 1.42 mm to -0.10 mm) (76) were analyzed using a stereology-based probe for quantitative analysis. Within the boxes randomly selected for counting, cells were counted if >60% of their perikaryon was in the plane of focus. The study included a total of 7 FPI and 6 sham animals, with at least two slices counted per animal, and the results were averaged. All analyses were performed with the rater blinded to the condition of the mice to minimize bias. In addition to quantitative stereological analysis, semi-quantitative densitometry analysis of GFAP and Iba-1+ cells were performed using Image J software (ImageJ).

For quantifying striatal CINs, ChAT-eGFP mice (naïve, H_2_O + FPI, EtOH + FPI) were anesthetized, perfused, sectioned, and then imaged using a confocal laser-scanning microscope (Fluoroview-1200, Olympus). Image processing was performed using Imaris 8.3.1 (Bitplane, Zurich, Switzerland). All GFP+ cells in the striatum from bregma 2 mm to -2.25 mm were manually counted for all groups using Imaris 8.3.1. The counts were then compiled based on the three groups and statistically analyzed.

### Electrophysiology

Electrophysiological recordings were conducted as previously described (16, 17, 31, 33, 42, 69, 77, 78). Specifically, coronal sections containing the DMS (thickness: 250 µm) were prepared at a speed of 0.14 mm/s in an ice-cold cutting solution containing (in mM) 40 NaCl, 148.5 sucrose, 4.5 KCl, 1.25 NaH_2_PO_4_, 25 NaHCO_3_, 0.5 CaCl_2_, 7 MgSO_4_, 10 dextrose, 1 sodium ascorbate, 3 myo inositol, and 3 sodium pyruvate, continuously oxygenated with 95% O_2_ and 5% CO_2_. After cutting, slices were incubated for 45 min in a 1:1 mixture of cutting and external solution containing (in mM) 125 NaCl, 4.5 KCl, 2.5 CaCl_2_, 1.3 MgSO_4_, 1.25 NaH_2_PO_4_, 25 NaHCO_3_, 15 sucrose and 15 glucose, and saturated with 95% O_2_ and 5% CO_2_.

The potassium-based intracellular solution was used for all cell-attached and current-clamp recordings in Figures 5c-f and 6d-e; this contained (in mM) 123 potassium gluconate, 10 HEPES, 0.2 EGTA, 8 NaCl, 2 MgATP and 0.3 NaGTP, with the pH adjusted to 7.3 using KOH. Cesium intracellular solution was used for all other whole-cell recordings; this contained (in mM) 119 CsMeSO_4_, 8 TEA.Cl, 15 HEPES, 0.6 ethylene glycol tetraacetic acid (EGTA), 0.3 Na_3_GTP, 4 MgATP, 5 QX-314.Cl and 7 phosphocreatine, with the pH adjusted to 7.3 using CsOH. The recording bath temperature was maintained at 32°C, with a perfusion speed set to 2-3 mL/min.

To inhibit glutamatergic transmission for recordings of spontaneous inhibitory postsynaptic currents (sIPSC), NBQX (10 µm) and AP5 (50 µm) were employed. Recordings of spontaneous excitatory postsynaptic currents (sEPSCs) were conducted by blocking all GABAergic transmission using picrotoxin (100 µm). CINs in slices were identified based on color and were held at -60 mV. Spontaneous cell-attached CIN firing activity was recorded for 5 min to calculate the average firing frequency. sIPSCs and sEPSCs were recorded for 3 min to determine average frequency and amplitude. The sag was calculated at a current injection of -400 pA. For measuring AMPA-induced currents in CINs, AMPA (5 µM) was applied via bath perfusion for 1 min, with holding currents recorded at a frequency of 0.2 Hz (30, 31, 79).

### Live Tissue Confocal

Acute brain slices were imaged in a custom-made chamber using an Olympus FluoView FV3000 confocal microscope, with flowing aCSF and saturated with 95% O2 and 5% CO2. The microscope was equipped with a 10x NA 0.3 and a 40x NA 0.8 water immersion objective, and 488 nm and 561 nm lasers. The imaging sample rate was 2-3 frames per second. Imaging parameters, including laser intensity, HV, gain, offset, and aperture diameter, were consistently maintained across sessions. MATLAB scripts for data analysis are available upon request.

### PLX 5622 Preparation

To prepare the 5 mg/mL (12.65 mM) solution of PLX 5622 (CHEMGOOD, C-1521), it was initially dissolved in dimethyl sulfoxide (DMSO, Sigma, D8418) to create a DMSO-PLX5622 stocking solution. The solvent for the DMSO-PLX5622 was 20% ethoxylated hydrogenated castor oil (MedChemExpress, HY-126403) in saline. The final solution involved diluting the DMSO-PLX5622 stocking solution in the 20% ethoxylated hydrogenated castor oil in saline solvent, resulting in a mixture of 5% DMSO-PLX5622 and 95% 20% ethoxylated hydrogenated castor oil in saline. The mixture was ultrasonicated until clear.

### Statistics

Data were analyzed using a Student’s t-test, one-way analysis of variance (ANOVA), or two-way ANOVA with repeated measures (two-way RM ANOVA), followed by the Tukey post hoc test. Significant differences were determined at *p* < 0.05 with a 95% confidence interval. Data is plotted as box and whisker plots, representing 25^th^ and 75^th^ percentile and min/max respectively. The horizontal line on the box plot represents the median. Analysis was conducted using SigmaPlot 12.5 software.

### Study Approval

All animal care and experimental procedures were approved by the Texas A&M University Institutional Animal Care and Use Committee.

## Data availability

Data will be made available from the corresponding author upon request.

## AUTHOR CONTRIBUTIONS

H.G. designed experiments, performed histology, behavioral, electrophysiological experiments, analyzed data, and wrote the manuscript; J.I. performed FPI surgeries, histology, behavioral experiments, analyzed data and revised the manuscript; Y.H. and W.D. performed behavioral experiments; W.D. wrote the early draft of the manuscript; S.M. performed FPI surgeries; R.C. and W.P. performed confocal imaging; R.C. generated MATLAB scripts and analyzed data; X.W. performed animal breeding; X.X., A.R. and G.J. performed data analysis; K.M. and K.O. assisted with behavior and histology; V.V. revised the manuscript; L.S and J.W. designed experiments, supervised the project, and wrote the manuscript. We also thank Gabriella Hollingsworth for assistance with behavior and histology.

