## Supplementary Figures 1-2 for "Traumatic Brain Injury Exacerbates Alcohol Consumption and Neuroinflammation with Decline in Cognition and Cholinergic Activity"

**Supplementary Figure 1.**

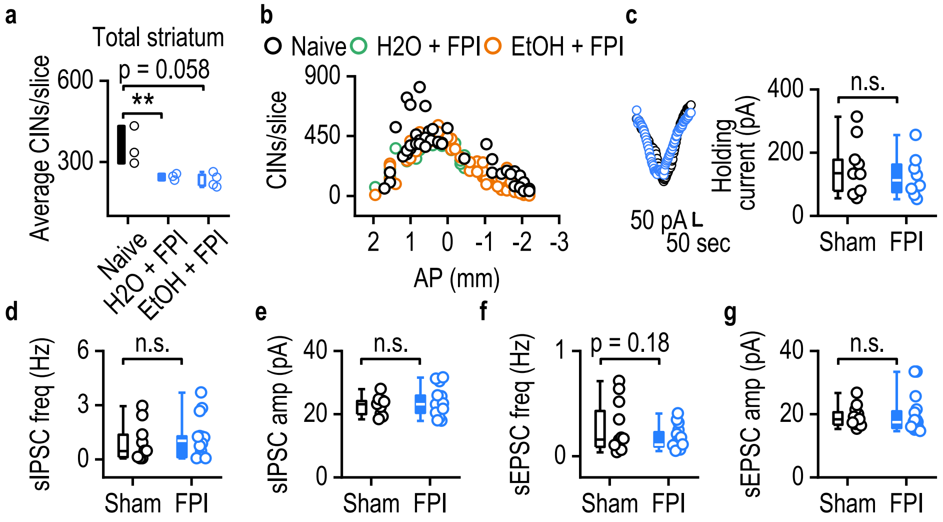


**a,** The average number of CINs per slice was reduced in both FPI and FPI with alcohol groups compared to naïve controls. **b,** Distribution of CINs per slice relative to the distance from bregma in naïve, FPI, and FPI with alcohol groups. **c-g,** There were no significant differences between the sham and FPI groups in DMS CINs in terms of AMPA-induced currents (c), spontaneous inhibitory current (IPSC) frequency (d), sIPSC amplitude (e), spontaneous excitatory current (sEPSC) frequency (f), and sEPSC amplitude (g). n.s. (not significant; c-g), ***p* < 0.01. Unpaired t-test (a, c, d, e, f, g). n = 3 mice (a, b, Naive), 4 (a, b, $H_2O$+FPI), 3 (a, b, EtOH+FPI). n = 10 neurons from 3 mice (10/3, c, sham), 8/3 (c, FPI), 12/2 (d, e, sham), 16/3 (d, e, FPI), 9/2 (f, g, sham), 14/3 (f, g, FPI).

**Supplementary Figure 2.**

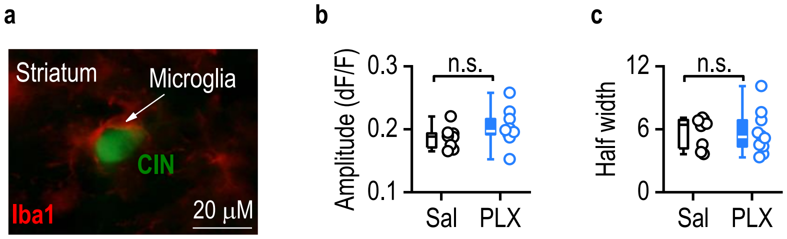


**a,** Iba-1 staining revealed a close spatial association between microglial cells and cholinergic interneurons in the striatum. **b-c,** There were no changes in the amplitude (b) or width (c) of acetylcholine release in the striatum after PLX administration. n.s. (not significant; b-c). Unpaired t-test (b, c). 8/4 (b, Sal), 10/4 (b, PLX), 9/4 (c, Sal, PLX).
